# HiCPlus: Resolution Enhancement of Hi-C interaction heatmap

**DOI:** 10.1101/112631

**Authors:** Yan Zhang, Lin An, Ming Hu, Jijun Tang, Feng Yue

**Affiliations:** Department of Computer Science and Engineering, University of South Carolina, Columbia, SC, USA; Bioinformatics and Genomics Program, Huck Institutes of the Life Sciences, Penn State University, University Park, PA 16802, USA; Department of Quantitative Health Sciences, Lerner Research Institute, Cleveland Clinic Foundation, Cleveland, OH 44195, USA; Tianjin Key Laboratory of Cognitive Computing and Application, Tianjin University, Tianjing, China; Department of Biochemistry and Molecular Biology, Penn State School of Medicine, Hershey, PA 17033, USA

**Author notes:** Equal contributors (Yan Zhang, Lin An).

## Abstract

**Motivation:** The Hi-C technology has become an efficient tool to measure the spatial organization of the genome. With the recent advance of 1Kb resolution Hi-C experiment, some of the essential regulatory features have been uncovered. However, most available Hi-C datasets are in coarse-resolution due to the extremely high cost for generating high-resolution data. Therefore, a computational method to maximum the usage of the current available Hi-C data is urgently desired.

**Results:** Inspired by the super-resolution image technique, we develop a computational approach to impute the high-resolution Hi-C data from low-resolution Hi-C data using the deep convolutional neural network. We hypothesize that the Hi-C interaction heatmap contains the repeating features, and develop an end-to-end framework to map these features from low-resolution Hi-C heatmap to high-resolution Hi-C heatmap at the feature level. Our approach successfully reconstructs the high-resolution Hi-C interaction map from the low-resolution counterpart, which also proves that the Hi-C interaction matrix is a combination of the regional features. Besides, our approach is highly expandable, and we can also increase prediction accuracy by incorporating ChIA-PET data.

**Availability:** Source code is publicly available at https://github.com/zhangyan32/HiCPlus

## 1 Introduction

The high-throughput chromosome conformation capture (Hi-C) technique (Lieberman-Aiden et al., 2009) has become an effective tool to measure the spatial organization of chromosomes by providing the pairwise interaction frequencies across the entire genome. In the past several years, Hi-C technique significantly expands our vision of the chromosome spatial organization (Lieberman-Aiden et al., 2009;Dixon et al., 2012;Rao et al., 2014) and gene regulation machinery (Schmitt et al., 2016). The Hi-C technique is a combination of proximity ligation of DNA and high throughput pair-end sequencing technique. Thus, sequencing depth is a crucial factor in determining the quality of the Hi-C interaction map. Meanwhile, the sequencing depth determined resolution is one of the most important metrics to describe the quality of Hi-C interaction map, which is described as fixed genomic size of non-overlapping windows along the genome (Schmitt et al., 2016). The bin size is inversely proportional to the resolution.

Although no quantitative relationship between the number of the reads and the resolution is determined, the linear increase of resolution requires a quadratic increase in the total number of the sequencing reads (Schmitt *et al.,* 2016). Consequently, it is very expensive to obtain the high-resolution Hi-C interaction data due to high sequencing cost. Currently, many datasets available can call large-scale patterns such as A/B compartment(Lieberman-Aiden *et al.,* 2009) and Topologically Associating Domains (TADs) (Dixon *et al.,* 2012, 2015; Jin *et al.,* 2013), but cannot be used to identify the cis-element interactions in a refined scale. Therefore, it is urgent to develop a computational approach to take full advantage of the currently available Hi-C datasets to generate higher resolution Hi-C interaction heatmap.

For the Hi-C map, the resolution is carefully determined by a series of the factors (e.g. distribution of cutter sites and sequencing depth). Simply mapping the same set of the sequencing reads to the smaller bin size can generate a "high-resolution" Hi-C interaction heatmap but also lead to very sparse data, and very low signal-to-noise ratio. From such heatmap, we cannot tell biologically meaningful interactions from random collisions. Therefore, in our work, we start from low-resolution Hi-C interaction heatmaps that are proven reliable.

Deep Convolutional Neural Network (ConvNet) (LeCun et al., 1998, 2015) is a type of multi-layer perceptron structure inspired by the organization of the animal visual cortex (Fukushima, 1980;Serre *et al.,* 2007;LeCun *et al.,* 1998), and achieves great success in computer vision and natural language processing (LeCun *et al.,* 2015). In computation biology and bioinformatics, ConvNet has been employed to predict the functional targets of DNA sequence (Zhou and Troyanskaya, 2015;Alipanahi *et al.,* 2015;Kelley *et al.,* 2016;Zeng *et al.,* 2016;Quang and Xie, 2016) as well as the epigenetic state (e.g. methylations and gene expression level) from experimental assays (Singh *et al.,* 2016;Angermueller *et al.,* 2016).

In this work, we propose HiCPlus, which to our knowledge is the first approach to generate the high-resolution Hi-C interaction maps from low-resolution interaction maps. Our approach is inspired by the most recent advancements (Glasner *et al.,* 2009;Yang *et al.,* 2008;Dong *et al.,* 2016) in the single image super-resolution, which trains a model on a large database containing both high-resolution images and corresponding low-resolution images to reflect the relationship between the patches of high-resolution and the patches low-resolution images. When low-resolution images are presented, the trained model then converts them to high-resolution images.

In HiCPlus, we employ deep ConvNet can be used as an end-to-end model from low-resolution patches to high-resolution patches, which is easy to train without intermediate step (Dong *et al.,* 2016). The key hypothesis for HiCPlus is that the chromatin interactions form repeating regional patterns, and that each interaction is affected by the strength of adjacent interactions regardless of cell-types. Therefore, we propose that HiCPlus can be used to enhance the resolution of Hi-C by learning the regional interaction patterns on one cell-type where the high-resolution Hi-C data is already available and published and apply it to predict the high-resolution interaction of other cell-types with low-resolution Hi-C data.

## 2 Methods

### 3.1 Overview

In the resolution enhancement process, we use low-resolution Hi-C interaction map as input, to predict high-resolution Hi-C interaction map. In Fig. 1, we illustrate the overall framework of HiCPlus. For each interaction (pixel in heatmap), the information for prediction obtained from its neighboring regions on the corresponding low-resolution matrix as shown in Fig. 1(a). Fig. 1(b) shows the conceptual view of the ConvNet. We will discuss the detail in later section. In Fig. 1(c), we present the pipeline of the HiCPlus. Instead of predicting every single interaction on the high-resolution matrix, we predict a block of interactions on the high-resolution matrix to accelerate the training speed.

(1) Divide Hi-C matrix into multiple square-like sub-regions with fixed size, and each sub-region is treated as one sample. For example, each sub-region is about 1Mb × 1Mb which contains 100 × 100 = 10,000 pixels at 10Kb resolution and 25 × 25 = 625 pixels at 40Kb resolution. We only investigate the regions where the genomic distance is less than 1Mb. To avoid the scenario in which the model is over-sensitive to the high interaction value, we set an upper threshold (100) for the interaction intensities.
(2) Interpolate the low-resolution matrix to the size of the high-resolution matrix. Interpolation can also be regarded as a simple method for the resolution enhancement, and common interpolation method includes simply scale, bilinear interpolation and bicubic interpolation. In this work, we use bicubic as our pre-processing strategy. Although the matrix size after interpolation is the same as high-resolution matrix, we still call them low-resolution samples.
(3) The deep ConvNet is trained to learn the relationship between the low-resolution samples (after interpolation) and high-resolution samples in the training stage, and predicts the high-resolution samples from low-resolution samples.
(4) The predicted high-resolution samples are recombined to the entire Hi-C interaction matrix. Considering the samples have a surrounding padding region which is removed during the prediction by ConvNet, the proper overlap is necessary when dividing the Hi-C interaction matrix to the samples in the first step.

**Figure 1.**
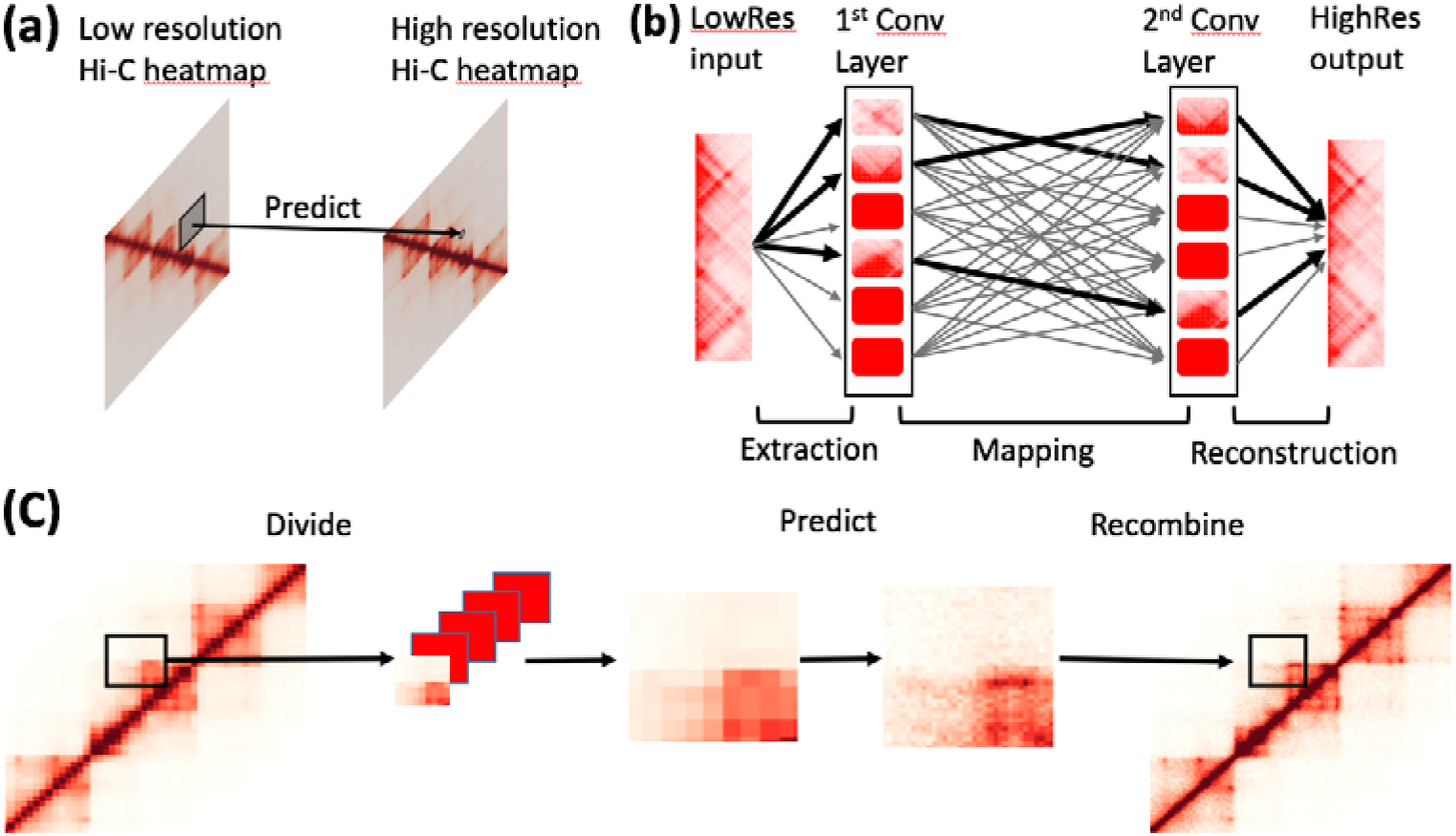
Illustration of the HiCPlus pipeline. (a) HiCPlus leverage information from nearby interactions to estimate contact frequency in certain location. (b) Conceptual view of the network structure in HiCPlus; (c) Schematic diagram for the HiCPlus model: regional interaction features (e.g. loops, domain borders) are learned from the low resolution matrix with available the high resolution Hi-C matrix as the training set. And predictions are made in the interpolated low resolution Hi-C matrix.

### 3.2 ConvNet structure

For the ConvNet, the input is a vector of *m × N × N* interpolated low-resolution samples, and the output is a vector of predicted high-resolution samples with the same dimensions, where *m* is the number of samples and *N ×N* is the size of each sample. In the training stage, the high-resolution Hi-C samples are also provided in the same format. We denote the ConvNet model as *F*, the low-resolution input as *X*, the predicted high-resolution output as *Y*, and the experimental high-resolution Hi-C as *Y'(Y* is also regarded as ground truth in this section). Mean Square Error (MSE) is used as loss function in the training process. Therefore, the goal of the training process is to generate F with minimized the MSE.

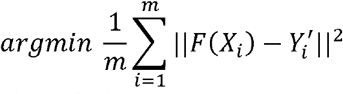

As shown in Fig. 1(b), the ConvNet in HiCPlus has three layers, serving for extracting and representing patterns on the low-resolution matrix, non-linearly mapping the patterns on the low-resolution matrix to high-resolution matrix, and combining the high-resolution pattern to generate the predicted matrix, respectively. Here we discuss each layer in detail.

#### 3.1.1 Pattern extraction and representation

In this stage, the input is the interpolated low-resolution *f*_1_ × *f_1_* size matrix, and the output is generated for the next layer by the formula

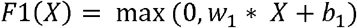

where * denotes the convolutional operation, *X* is the input matrix, *b_1_*is the bias, and *w_1_* is *n_1_* × *f_1_* × *f_1_* matrix. The *n_1_* and *f_1_* are the filter numbers and filter size, respectively. Both *n_1_* and f are hyperparameters in the ConvNet, and so we set *n_1_* to 16 and *f_1_* to 5 (the HiCPlus is not sensitive to these hyperparameters). The Rectified Linear Unit (ReLU) (Nair and Hinton, 2010) is utilized as the non-linear activation function.

We extract the patterns by convolving the input matrix by n1 number of filters where each filter represents a local pattern. The patterns may include interaction peaks, the right-angle of the TADs, the edge of TADs, etc., but the model may extract such pattern in a more abstract and complicated way which is beyond our knowledge.

#### 3.1.2 Non-linear mapping between the patterns on high-resolution and low-resolution maps

This stage is shown as the middle part of the Fig. 1(b), where the patterns on the low-resolution matrix are mapped non-linearly with the patterns on high-resolution matrix using the formula:

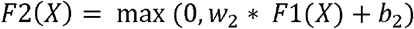

where *F*_1_(*X*) is the output from the previous layer, *b_2_* is the bias, and *w_2_* is *n_2_* number of *f_2_* × *f_2_* matrix. In this layer, we set *n*_2_ to 16 and *f_2_* to 1 as it is a process of non-linear mapping.

#### 3.1.3 Combining the high-resolution patterns to generate the predicted high-resolution maps

We employ the following formula to generate the predicted high-resolution Hi-C matrix from the results of the second layer

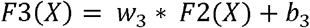

where *F2*(*X*) is the output from the previous layer, *b_3_* is the bias, and *w_3_* is *n_3_* number of *f_3_* × *f_3_* matrix. In this step, the non-linear activation function is not required, and the filter number *n_3_* is set to 1 to generate the predicted results.

Overall, the *F* has the parameters *Θ*= {*w*_1;_ *w*_2_, *w*_3_, *b*_1_*, b_2_*, *b*_3_}. The goal of the training process to obtain the optimal *Θ*to minimize MSE on the samples in the training set.

## 3 Results

### 3.1 Single interactions are predictable from regional interaction patterns

As shown in Fig. 1(a), we hypothesize that the interaction intensity for each interaction is related to its local neighborhood region. If this hypothesis stands, the read count of an interaction in the Hi-C matrix could be predicted from reading counts in its nearby region. To test this hypothesis, we systematically selected interactions formed by two loci are less than 0.6Mb (60 of 10Kb bins) away. The training sets are from Chr1-17 on GM12878 cells, and the testing sets are from Chr18-22 in the same cell type (Rao *et al.,* 2014). Then, we predicted the values of these interactions from nearby regions by using ConvNet to test: 1) the value is predictable, and 2) the ConvNet is an effective method to be employed as the prediction approach. We set two benchmark methods for the comparison: 1) use the average value of the distance as predicted value; 2) use the simple average value of the nearby 4 interactions or 8 interactions as the predicted value. We predicted the value of the interactions by multiple methods and list the result in Fig. 2. We first used the average at the distance as the predicted value and the MSE is higher than any imputation method. The Pearson correlation and Spearman rank correlation cannot evaluate the results from this method since the variance of predicted value is zero.

**Figure 2.**
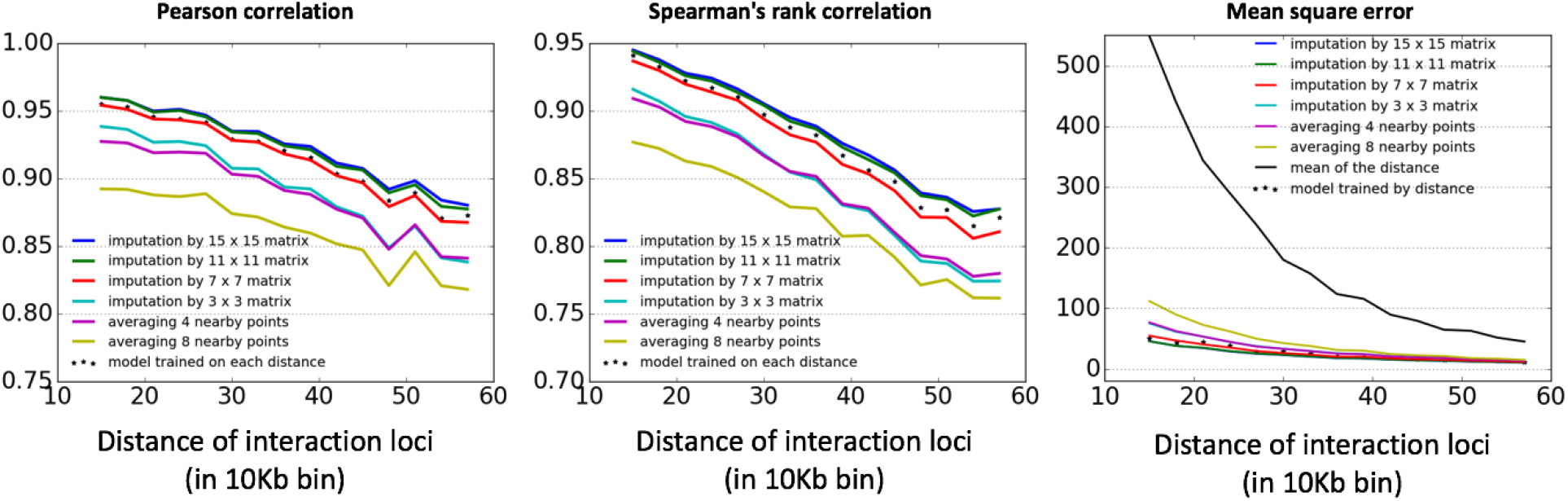
Model performance on single point imputation from the nearby points. We tested the ConvNet with multiple sizes of neighborhood interactions as well as the naive imputation approaches. We also train the separated models for the points at each distance as shown in asterisk makers. The performance was evaluated with experiment high resolution Hi-C matrix in each distance.

An alternative method to impute the value of an interaction is using the average value of its nearby interactions, including 4 interactions average (up, below, left and right) and 8 interactions average (up, below, left, right, up-left, up-right, below-left and below-right). The averaging method can obtain the values which are close to the true values (both Pearson correlation and Spearman rank correlation is larger than 0.7 at each distance) and MSE value is less than the averaging value of entire distance. As shown in the Fig. 2, the result of a direct average at a larger scale (8 interactions) has higher MSE and lower correlation than small scale (4 interactions). It indicates that the interactions closer to the target interaction are more important than other interactions in the model. To exploit the information from the larger scale nearby matrix, we applied the ConvNet, which has the similar network structure with HiCPlus, to impute the value of the target point. The results from ConvNet outperform simple averaging nearby points at every case.

We also employed different sizes of the matrix to investigate the optimal range of the nearby points to be used. Four models are trained with (3 × 3), (7 × 7), (11 × 11), and (15 × 15) matrix of the nearby region. As we expected, the larger nearby region yields better performance. We also observed that imputation with (15 × 15) matrix is nearly the same as using (11 × 11) matrix, indicating the points further than (11 + 1) / 2 = 6 has merely no contribution to the imputation as shown in Fig. 2. To confirm the result, we trained the model using (15 × 15) matrix with extra epochs, and no further improvement is observed.

It is also important to compare the performance of training one single model for samples in all distances with training separated models for each distance. In Fig. 2, we use asterisk mark to show the predicted results from separated models, which is also trained on (11 × 11) matrix. The performance of separated models decreases comparing with one single model, indicating that the points at different distance share very similar pattern. Training separated models at different distance dramatically reduces the number of the samples in the training data sets, which explains the decreases of model performance.

### 3.2 Imputation result from the HiCPlus

In this section, we used the normalized 10Kb resolution Hi-C interaction matrices (access code GSE63525) (Rao *et al.,* 2014). The training data sets were obtained from Chr1 on K562 cells, and the testing data sets were obtained from Chr18 on GM12878 cells. The low-resolution matrices (with reduced size and before the interpolation) were generated by merging the nearby high-resolution interactions and the mean value of the combined interactions are used as the new value. The samples were generated by systematically dividing the entire Hi-C matrix into 0.4Mb × 0.4Mb base pair regions with 0.15Mb overlap at each direction. We only investigated the region where two loci are within a genomic distance of 1.2Mb (Expanding the genomic distance range has the same accuracy but reduces the training speed significantly).

In Fig. 3(a), we pick a region (center at chr18: 2.95Mb - 3.20Mb) which contains several obvious patterns to illustrate the performance of HiCPlus, and we also plot the output from two interpolation-based methods. By the visual examination, we noticed that, comparing with bicubic interpolation and bilinear interpolation, HiCPlus enhanced the loops at the correct location and sharpened the edge of the domain. Overall, HiCPlus successfully reconstructed the real high-resolution Hi-C data comparing with the other interpolation methods.

**Figure 3.**
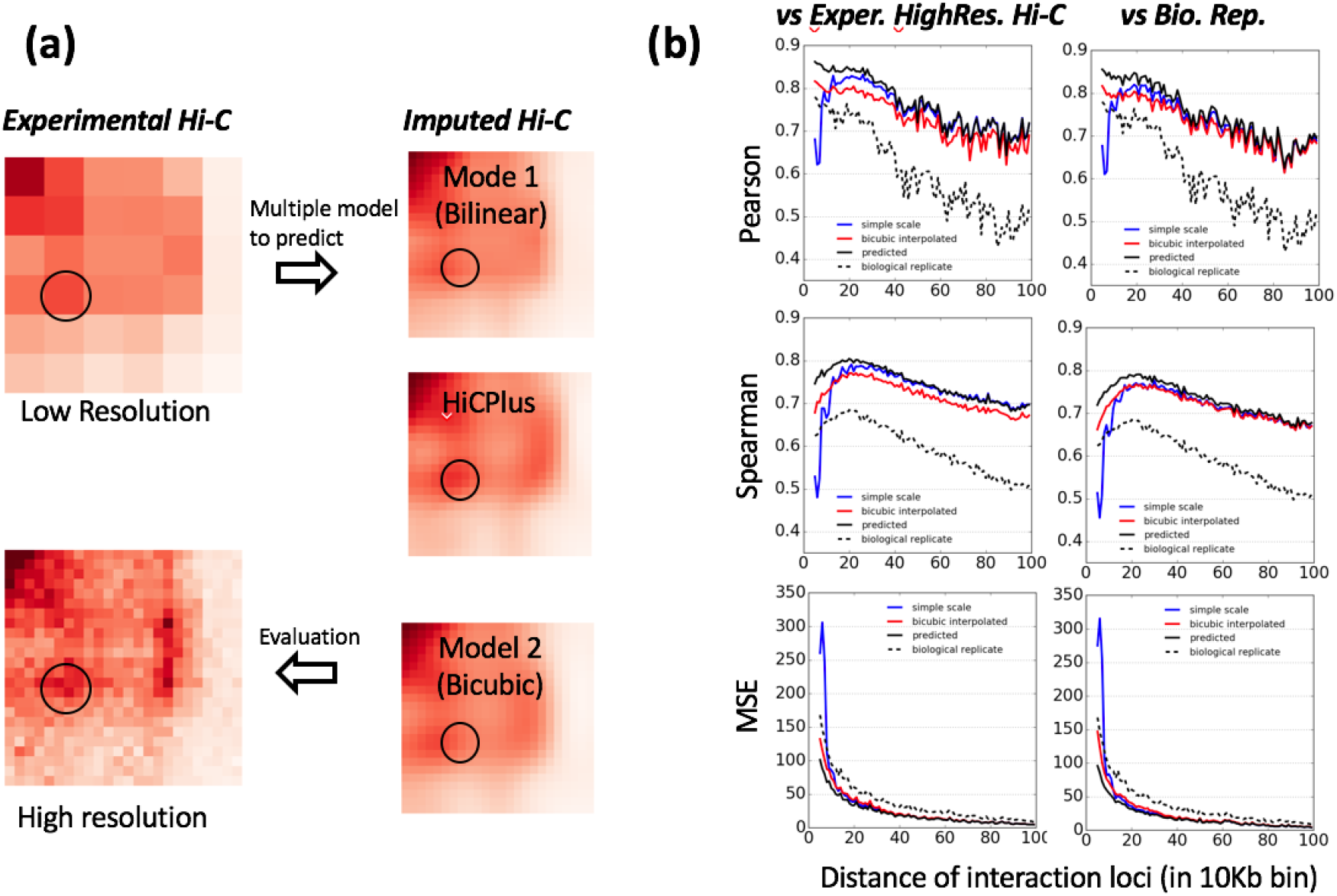
Imputed high resolution contacts are close to experiment data. (a) Comparison of HiCPlus with other models. HiCPlus results captured location of high frequent contacts (circle region) in the refined resolution. (b) HiCPlus predictions agreed with ChIA-PET data (blue dots). (c) Quantitative comparison of prediction at each genomic distance.

To quantitatively evaluate the performance of HiCPlus, we calculated the MSE, Pearson correlation and Spearman rank correlation at each genomic distance (Fig. 3(b)). We compared the imputation from HiCPlus with bicubic interpolations, simple scale and biological replication. The result from bilinear interpolations was worse than bicubic interpolation, so it was not included in the plot. As shown in Fig. 3(b), HiCPlus achieved the highest correlation and the lowest MSE at all genomic distances. For the region where two loci are close together, the bicubic was better than simple scale while at the regions two loci are far away, the simple scale is better. Noticeably, the biological replication had the least similarity with the original Hi-C matrix, comparing with any imputation methods. To further study the stability of the results, we also calculated the similarities between the imputation results with the biological replication, which was never posed to neither the training nor testing process. Surprisingly, the imputation matrix from HiCPlus, as well as bicubic interpolation, nearly maintained the same performance to the biological replication version of Hi-C while the simple scale approach had some performance drop but all of the imputations still had higher correlation than biological replication. One possible explanation is that the Hi-C data is derived from a large volume of cells and due to the systematic variability in the experiment, the noise level is high. For the low-resolution Hi-C matrix, the process of averaging the nearby interactions can be served as the de-noising process.

### 3.3 Validation of the results from HiCPlus

In the previous sections, we have shown HiCPlus outperformed all the existing methods for low-resolution to high-resolution imputation in several quantitative metrics such as MSE, Pearson correlation and Spearman rank correlation. In this section, we also demonstrate the biological significance of HiCPlus.

ConvNet has achieved great success in several domains (e.g. computer vision and natural language processing), but its internal mechanism is still largely unknown. To test whether the ConvNet is working properly, and study the region where ConvNet utilizes to make correct decision is of great importance. For example, in the dog breed classification problem, the most important region should be the dog face if the ConvNet is working correctly (Zeiler and Fergus, 2014). In our work, we expect HiCPlus to reconstruct the biologically meaningful patterns such as loops and TADs. Some other patterns are useful to increase the correlation value and reduce the MSE but are less biologically meaningful, such as the distance effect and the cutting site/mappability bias. Since the ConvNet in our model takes the bicubic interpolated matrix as input and predicts the high-resolution Hi-C matrix, we evaluated improvement (decrease in MSE or increase of the correlations versus high-resolution matrix) from input matrix to output of the ConvNet. For the region with known biological importance pattern (e.g. loops and TADs), the improvement should be larger than those without such patterns. In Fig. 4(a) and Table 1, we showed an example of comparison between two regions. In the upper sample of the Fig. 4(a), a pair of loop peaks exist while in the lower sample no biological important patterns exists. In Table 1, we compared the improvements of these two samples, and obviously, the sample with loop has significant improvement in all of three metrics. The observation indicates HiCPlus successfully reconstructed the biological important patterns.

**Figure 4.**
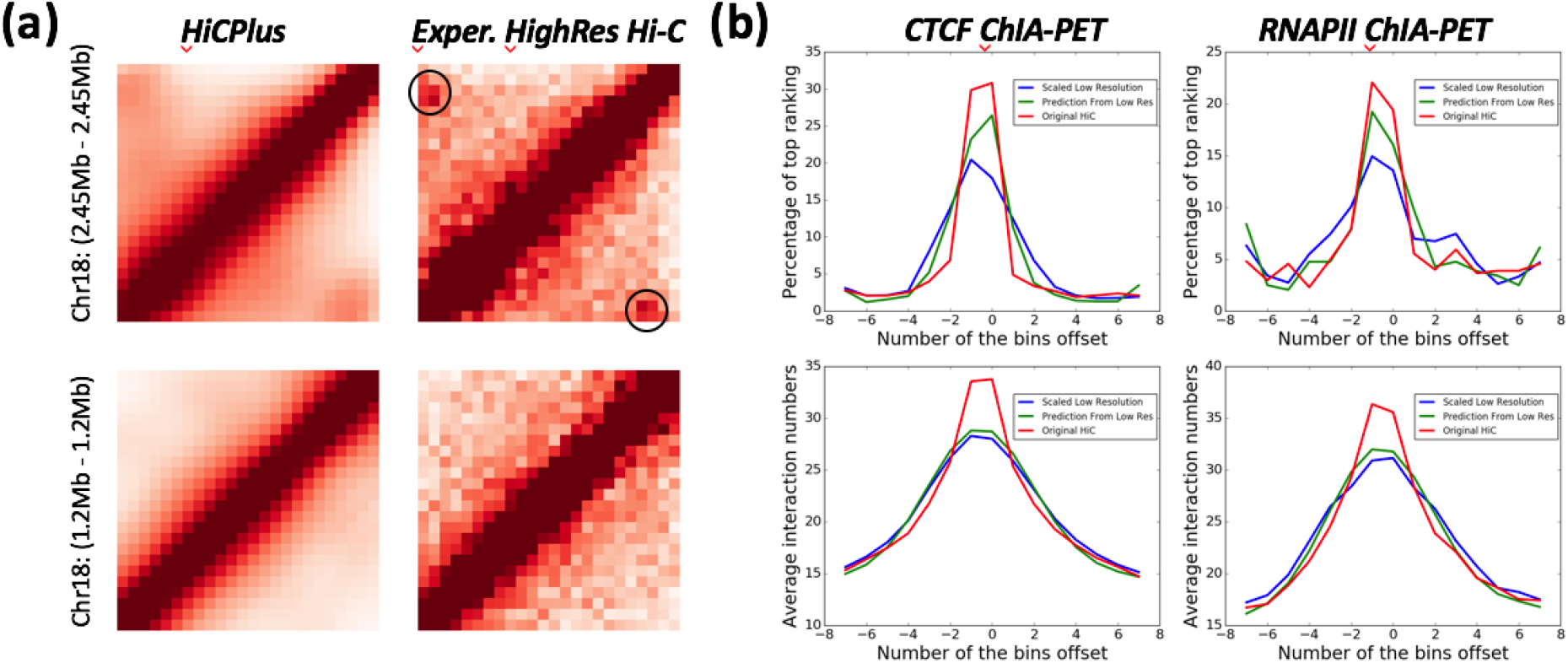
Validation of the imputation results. (A) We compared two regions: the top one contains a loop and TAD edge, and the lower one doesn’t contain any biologically meaningful pattern (circle region). We listed the improvement at Table 2. The upper has larger improvement comparing with bicubic interpolated matrix in all of the metrics; (B) We used the ChIA-PET interaction peak as the base point and investigated the window nearby. The upper two is the percentage of the top ranked point at each relative location, and the bottom two are the average interaction intensities at the relative location with the ChlA-PET peaks.

**Table 1.** Improvement of the ConvNet predictions at different regions *The table describes the samples in Fig 4(a). All metrics are calculated versus experimental high-resolution matrix

|  | Chr 18: 1.2Mb - 1.2Mb |  |  | Chr 18: 2.45Mb - 2.45Mb |  |  |
| --- | --- | --- | --- | --- | --- | --- |
|  | MSE | Pearson | Spearman | MSE | Pearson | Spearman |
| Bicubic | 87 | 0.950 | 0.936 | 78 | 0.959 | 0.926 |
| Predicted | 49 | 0.969 | 0.955 | 60 | 0.967 | 0.932 |
| Change | -38 | 0.019 | 0.018 | -18 | 0.008 | 0.006 |

Besides investigating the importance of patterns, we also validated the prediction with publicly available ChIA-PET data, which is an alternative approach to investigate the spatial organization of the genome (Fullwood *et al.,* 2009;Li *et al.,* 2010;Tang *et al.,* 2015). Similar to Hi-C technique, ChIA-PET also measures the interaction intensities between loci on the genome, but unlike Hi-C unbiased covers genome-wide interactions, ChIA-PET only captures the interactions between a pair of loci that bind with certain protein. Since the ChIA-PET data sets are produced by an independent process, we employed the ChIA-PET data (GSE72816) (Tang *et al.,* 2015) to validate our imputation results. The ChIA-PET interactions (e.g. CTCF-binding sites) are comparable to the chromatin loop peaks identified from high resolution Hi-C data, so we decided to use the ChIA-PET data to as an independent data source to validate our results. Since the low-resolution Hi-C matrix is an average of the high-resolution Hi-C, we compared the interaction intensity values of locations where ChIA-PET peaks observed on the high-resolution matrix with those on the low-resolution matrix. As shown in Table 2, for the real high-resolution Hi-C data, the interactions that overlap with ChIA-PET peaks have higher value than its neighbors, and the prediction from HiCPlus shows the similar trend.

**Table 2.**
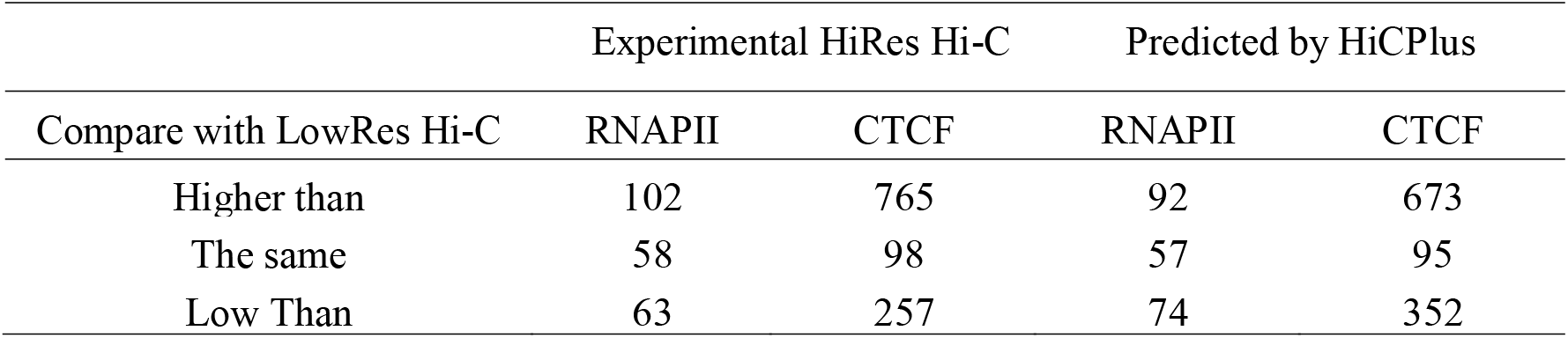
Comparison of the values with the low-resolution Hi-C matrix

Since the genomic distance between two interacting loci strongly affects chromatin interaction frequency in Hi-C data, we also compared each value with its upstream and downstream at the same distance. In Fig. 4(b), we chose all points with valid ChIA-PET interactions as the base points and calculated the average interaction frequencies and rankings within the window from −7 bins to +7 bins. In the upper figures of Fig. 4(b), we plotted the average interaction frequency around the ChIA-PET peaks. As expected, the interaction frequencies at the ChIA-PET peaks, for both CTCF and RNAPII, are much higher than the background. The predicted results from HiCPlus outperformed the low-resolution Hi-C in the enrichment, indicating that the predicted results correctly reflected the biological signals. In the lower part of Fig. 4(b), we plotted the ranking-based results, which are the percentage of interaction intensity in each window. The contrast is higher than the upper plotting, and in most of the cases, the highest interaction frequency in this window overlaps with the ChIA-PET peaks.

### 3.4 Expansion the scope of the application

From the Fig. 1(b) and the concept of our structure of ConvNet, the procedure can be summarized as 1) input pattern extraction, 2) mapping input pattern and output pattern, and 3) output pattern recombination. In current work, the input is the low-resolution Hi-C interaction matrix. Based on the concept of the ConvNet structure, we proposed that the input is not limited to the low-resolution Hi-C matrix. Other information sources can also be added into the network. As mentioned in the previous section, since ChIA-PET is similar to Hi-C and only acquires the interaction of a subset of the genome, the sequence depth requirement of ChIA-PET is much lower than Hi-C for the same level of resolution. Therefore, we employed the ChIA-PET data together with the low-resolution Hi-C data to impute the high-resolution Hi-C data. The pipeline, ConvNet structure and training procedure nearly remains the same, and we showed the results in Fig. 5. We used the CTCF ChIA-PET data and RNAPII ChIA-PET data from Chr1 on GM12878 cells and the low-resolution Hi-C data from the same region. In Fig. 5(a) and Fig. 5(b), we compared the predicted results from the low-resolution Hi-C data only and predicted result from the combination of the low-resolution Hi-C data and the ChIA-PET data, respectively. We plotted the real high-resolution Hi-C data in Fig. 5(c). By comparing the predicted results with the truth, the predicted matrix with ChIA-PET data showed more significant loop peaks and sharp TAD edges compared with the predicted matrix only with the low-resolution Hi-C data. The quantitative metrics (e.g. MSE, Pearson correlation and Spearman rank correlation) also showed similar results. To further investigate the effect of the ChIA-PET data in the prediction, we plotted a heatmap reflecting the difference between two predicted matrices on the Fig. 5(d). The locations with ChIA-PET peaks are enhanced, and more interestingly, the surrounding region is suppressed to sharpen the contrast between the significant interaction regions and background. To further validate our results, we used Fit-Hi-C (Ay et al., 2014) detected significant interactions (with FDR < 0.01) from all Hi-C matrices (including experimental and imputed) and investigated the overlap as shown in Fig. 5(e).

**Figure 5.**
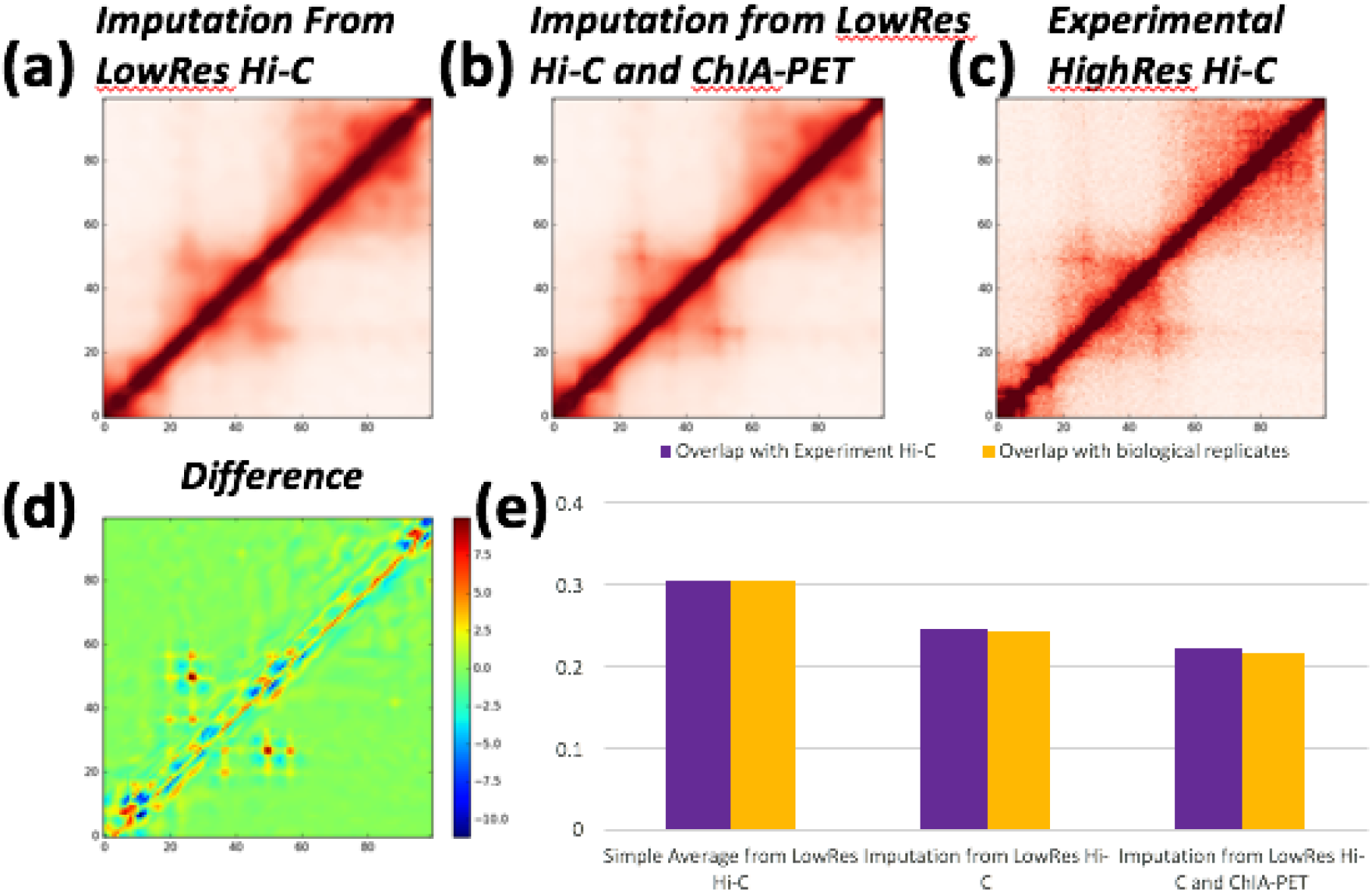
Expansion of the application. We used the two ChIA-PET matrices together with the low-resolution Hi-C matrix to predict the high-resolution Hi-C matrix. (a) The imputation from the low-resolution Hi-C matrix only; (b) The imputation from the low-resolution Hi-C matrix and ChIA-PET matrix; (c) The experimental high resolution Hi-C; (d) Difference between two predictions to illustrate the effect of the additional ChIA-PET data sets; (e) Validation from Fit-Hi-C. We plotted the percentage of the significant interactions generated from Fit-Hi-C which are NOT overlapped with the real Hi-C data.

In future, with more ChIA-PET data and similar data sets are available, additional data sets can be applied to the HiCPlus to obtain the better imputation results.

## Discussion and Conclusion

In summary, HiCPlus presents the first implementation of image scaling on computationally refining the resolution of Hi-C contact map. By leveraging information from adjacent interactions and learning regional patterns from K562 high-resolution Hi-C map, we improved the resolution of GM12878 Hi-C map from 40Kb to 10Kb. The refined Hi-C map agrees with interactions identified by high-resolution Hi-C and ChIA-PET experiments.

We showed that repeating local patterns could be learned from a group of local chromatin interactions in Hi-C data. Although not all regional patterns are directly related to any biological functions (e.g. distance effect and systematically bias in Hi-C experiment), some patterns learned by HiCPlus are associated with regulatory units (e.g. edges of TADs and loop anchors). The appearance of those regulatory units related pattern in our approach provides a powerful approach to use the deep ConvNet model to discover the pattern in a fully automatic way. One may train ConvNet in known loop regions and apply the trained model to the entire Hi-C matrix to search for possible loops. Deep ConvNet-based models have been widely used in recent studies, however, it is still difficult to interpret the learning complex. Although the ConvNet in HiCPlus achieves success in the resolution enhancement, the patterns extracted by the ConvNet are still difficult to interpret, and we will continue to study the patterns in the Hi-C interaction matrix in the near future.

The HiCPlus is also expandable to exploit more information besides the low-resolution Hi-C matrix. Nearly all interaction data can be incorporated with the low-resolution data to generate more accurate results. Currently, the ChIA-PET is only available in several cell-types with very limited types of targets. HiCPlus has great potential when more ChIA-PET data or similar data sets become publicly available.

Overall, HiCPlus is a useful tool to investigate chromatin organization in tissue/cell types where high-resolution Hi-C data is not available. It will also be a boost to the study of cis-regulatory network by scaling up the resolution of interaction heatmap.

## Acknowledgements

We are also grateful to the NVIDIA Corporation for donation of a TITAN X GPU card through a NVIDIA Hardware Grant.

## Funding

This work is supported by NIH grants U01CA200060 and R24DK106766 (to FY), U54KD107977 (to MH), and by NSF award #1161586 to YZ and JT.

